# Determination of the amount of insecticide picked up by mosquitoes from treated nets

**DOI:** 10.1101/2020.08.07.241224

**Authors:** Mojca Kristan, Jo Lines, Harparkash Kaur

**Affiliations:** Department of Disease Control, London School of Hygiene & Tropical Medicine, London, UK Jo Lines; Department of Clinical Research, London School of Hygiene & Tropical Medicine, Keppel Street, London, UK

**Keywords:** malaria, *Anopheles gambiae*, pyrethroids, deltamethrin detection, HPLC, type 2 pyrethroid colorimetric test

## Abstract

**Background:** Insecticides used in vector control mostly rely on vectors being exposed through contact with treated surfaces, yet little is known about the amount picked up by the insect. Measuring this amount is relevant not only for determining the actual doses that are lethal to the mosquito, but also for understanding effects on the physiology and vector competence of mosquitoes. Insecticides at sub-lethal doses can affect both parasites developing inside mosquitoes and mosquito microbiota, hence it is important to understand the processes by which parasites are exposed to insecticide inside the insect. These doses will inevitably depend on the amount of insecticide that mosquitoes pick up when they come into contact with treated nets.

**Methods:** Three to five days old non-blood fed female *Anopheles coluzzii* mosquitoes were exposed to a long-lasting insecticidal net (PermaNet 2.0 containing 55 mg/m^2^ deltamethrin), using a wire ball frame, for 0.5-5.0 minutes. Our in-house developed colorimetric test was used to visually detect the amount of deltamethrin on different parts of the mosquito (legs, heads, thoraxes, abdomens) following exposure to the net. The amount of insecticide picked up by mosquitoes from the net over a range of exposure times was measured using a high-performance liquid chromatography with diode array detection (HPLC-DAD).

**Results:** The colorimetric test, designed to only detect the type 2 pyrethroids (i.e deltamethrin, α-cypermethrin and λ-cyhalothrin) on fabrics (e.g. ITNs) and sprayed walls, was successfully used for the first time to detect deltamethrin on mosquitoes following exposure to the net. The confirmatory HPLC-DAD analysis determined that after 2 min exposure up to 12 ng of deltamethrin adhered to mosquitoes following exposure to PermaNet 2.0 (mean = 5.2 ng/mosquito, SE = 1.9) and that the final dose depends on the length of exposure time.

**Conclusions:** This study demonstrated the potential of a screening (type 2 pyrethroid colorimetric test) and a confirmatory test (HPLC-DAD) to determine the amount of insecticide that adheres to mosquitoes on contact with treated surfaces. This has implications for a precise lethal dose determination and detection of specific insecticide that causes the greatest mosquito mortality in circumstances where mixtures of insecticides may be used to maximise effectiveness of interventions.

## BACKGROUND

Recent declines in malaria incidence across sub-Saharan Africa (SSA) have largely been attributed to a scale-up of insecticide-based vector control interventions, such as insecticide-treated nets (ITNs) and indoor residual spraying (IRS) [1]. However, only a small number of insecticides can be utilised for these interventions and pyrethroids are currently the only insecticides used on all ITNs, either alone or in combination with a synergist piperonyl butoxide (PBO) or a non-pyrethroid insecticides such as chlorfenapyr and pyriproxyfen [2]. Pyrethroids were first approved for use in mosquito control by the World Health Organization (WHO) in the 1970s [3]. They are neurotoxins affecting the *para* voltage-gated sodium channels (VGSC) on the mosquito’s neurons [4, 5]. They work well on nets and sprayed surfaces because of their rapid knock-down effect, killing properties, and a long residual action [6].

*Anopheles* mosquitoes in SSA generally feed every 2-3 days, once per gonotrophic cycle [7]. They come into contact with pyrethroids with the tips of their legs when they rest on a sprayed wall after taking a blood meal or come into contact with an ITN while trying to blood feed [8]. There are several possible ways pyrethroids can enter the mosquito’s body [9]. It is not known precisely how these insecticides enter and reach their target site VGSCs [10], nor if they accumulate in tissues or are immediately metabolised, and whether their metabolites exhibit any insecticidal activity [11] or if they might be potentially sporontocidal [12, 13, 14].

Uptake of insecticides from treated nets or surfaces is variable depending on the formulation, active ingredient availability, contact time, knockdown, temperature and irritant effects. Insecticides for IRS are available as different formulations, which should provide long lasting residual effect and bioavailability on a number of different surfaces [2, 15], whereas insecticide is incorporated within the netting or bound around the net fibers in long-lasting insecticidal nets (LLINs) [16]. There is a lack of data about the amount of insecticide picked up by mosquitoes after they come into contact with a treated surface, either a net or a wall. Only a few studies have attempted to measure the amounts in the laboratory or field conditions, using gas chromatography to detect DDT or dieldrin, or more recently liquid chromatography coupled to high-resolution tandem mass spectrometry (UHPLC-MS/MS) to detect different pyrethroids and organophosphate pirimiphos-methyl [17, 18, 19].

Knockdown resistance (*kdr*) mechanism is associated with reduced irritant effects of pyrethroids, implying that resistant mosquitoes tend to search longer to feed, remain in contact with treated surfaces longer before taking off and acquire more insecticide through contact. This may result in a total dose picked up to be high enough to kill even homozygous *kdr* resistant mosquitoes [20, 21, 22]. On the other hand, mosquitoes with metabolic resistance mechanisms may detoxify the insecticides faster than susceptible mosquitoes and while this is advantageous for survival, resistance is often thought to be associated with fitness costs to the insect [23]. Developing new vector control tools, including the next-generation LLINs, will require a more thorough understanding of how they function in terms of their physiological mode of action and mosquito behavior around them [8]. The minimum duration of LLIN contact necessary to deliver an effective insecticide dose is not known, but mosquito-LLIN interactions have been described and average contact times measured (<2 min contact time for a single mosquito) [8, 24].

The degree to which the insecticide is lost from LLINs to make them ineffective and the length of their useful life can vary considerably [25]. Measuring the rate of insecticide loss or, in case of spraying, monitoring of insecticide residues reaching the surface, can provide valuable information to vector control programs. Various high-tech techniques, such as high performance liquid chromatography (HPLC) and gas chromatography (GC) are often used for measuring the amounts of pyrethroids but are reliant on sophisticated laboratory-based equipment and need both expertise and experience to utilize [26, 27, 28]. More recently, colorimetric tests have been developed and used for detection of pyrethroids [29, 30, 31, 32] and carbamates [33], as well as biosensors using glutathione-S-transferase for pyrethroids [34, 35, 36] and DDT [37], and DDT dipstick assays [38] on fabrics and IRS. To date these methods have not been used to detect insecticides on exposed mosquitoes.

The ability to determine the quantity of insecticide mosquitoes pick up on contact with treated surfaces will provide invaluable information regarding the active ingredient that causes the greatest mosquito mortality in circumstances where mixtures of insecticides are used. Overall, the findings will provide important information to assess and maximize the effectiveness of interventions. Our aim was therefore to test whether the type 2 pyrethroid colorimetric test developed by Kaur and Eggelte [29] can be used to detect the presence of deltamethrin on mosquitoes, to precisely measure the amount of insecticide mosquitoes pick up during contact with a LLIN and to compare the results of the screening (type 2 pyrethroid colorimetric) and confirmatory (HPLC-DAD) analyses.

## METHODS

### Mosquito insecticide exposure

Three to five days old non-blood fed female *Anopheles coluzzii* mosquitoes (susceptible N’gousso strain [39]) were exposed in groups of 10 insects to PermaNet 2.0 (55 mg/m^2^ deltamethrin), using a wire ball frame [40]. For HPLC-DAD and associated colorimetric test, mosquito exposure times varied between 0.5-3 minutes, in 0.5-minute increments (Fig. 1). Mosquitoes were exposed to the net for 5 minutes to confirm whether detection of deltamethrin on whole mosquitoes and different body parts is possible using type 2 pyrethroid colorimetric test, (Fig 1).

**Fig. 1.**
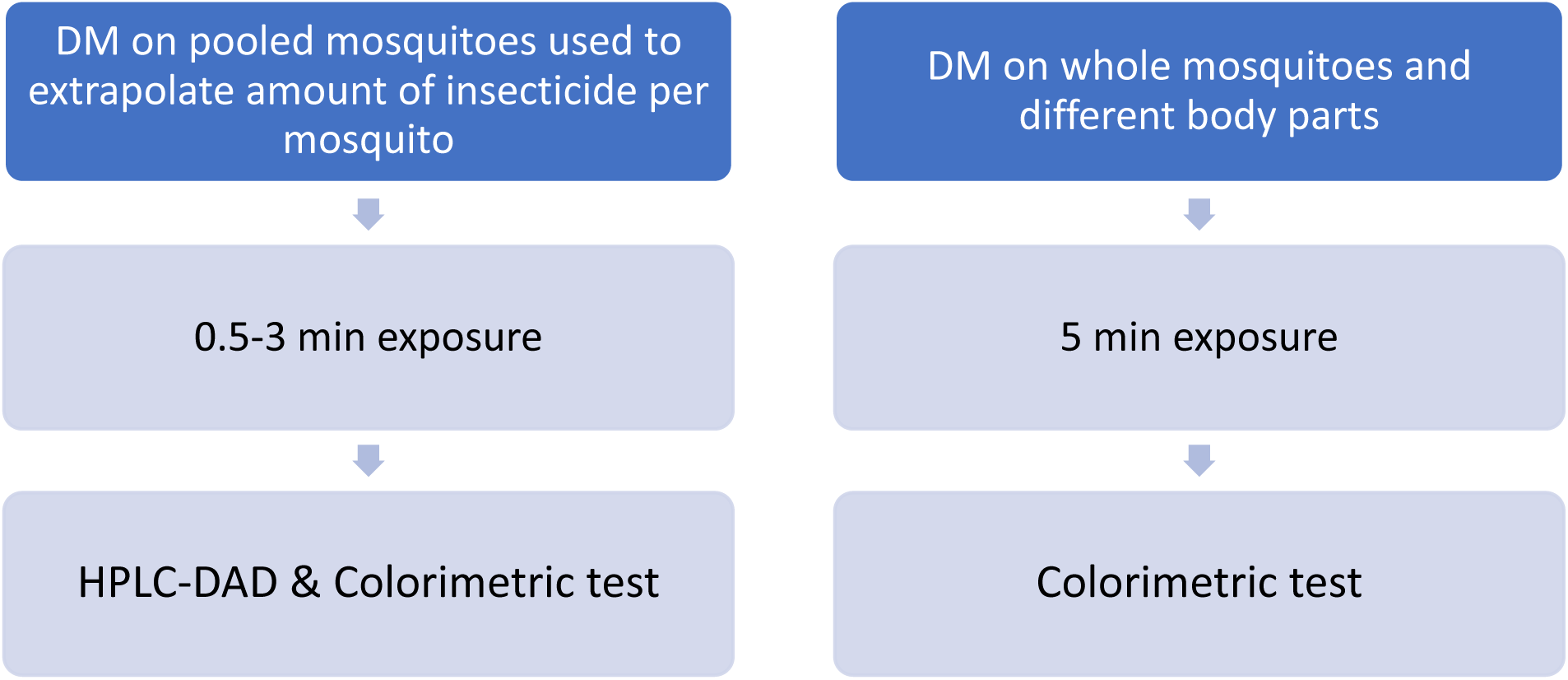
A diagram showing a summary of two sets of experiments of mosquitoes exposed to LLINs (DM = deltamethrin).

Following insecticide exposure, the mosquitoes, together with unexposed controls, were killed by freezing within 5 minutes (to prevent the enzymes from degrading the insecticides) and were stored at −20°C until screening and confirmatory chromatographic analysis.

### High Performance Liquid Chromatography with diode array detection (HPLC-DAD)

#### a) Mosquito sample preparation

The exposed and control mosquitoes were placed in Eppendorf tubes in pools of 10, relating to their exposure time (Table 1).

**Table 1.**
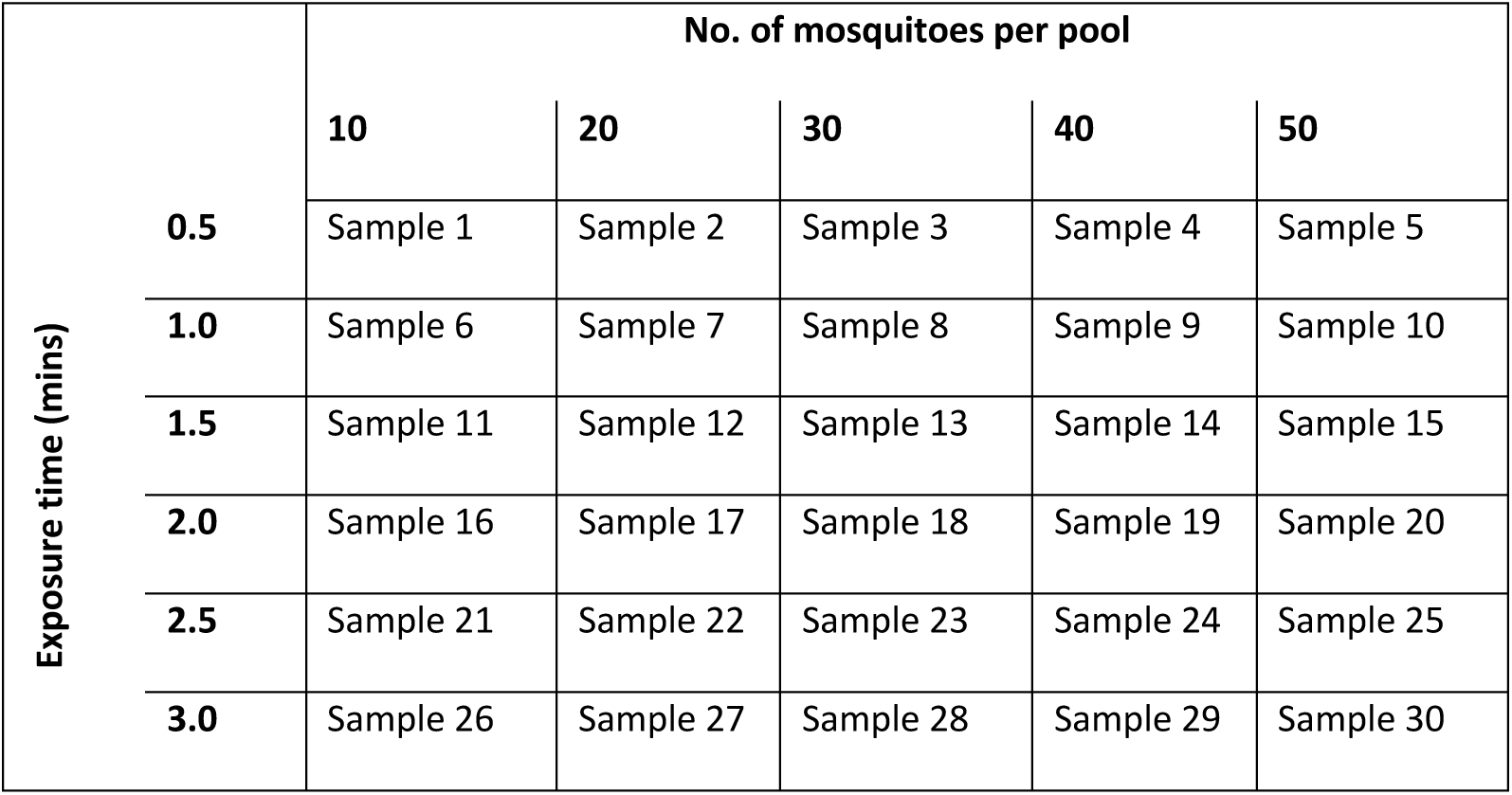
Details of exposure times and the number of mosquitoes pooled in each sample used for HPLC-DAD and type 2 pyrethroid colorimetric test experiments (controls not shown in this table).

400 µl of acetonitrile (CH_3_CN) were added to each sample. Mosquitoes were first ground roughly using plastic pestles, then sonicated for at least 20 minutes or homogenized using a Qiagen Tissue Lyser II (Qiagen, Hilden, Germany) with 3 mm stainless steel beads, making sure all mosquitoes in each sample were completely crushed. Samples were then vortexed for 10 seconds and centrifuged for 4 minutes. Supernatant was removed into new tubes and used for HPLC-DAD and type 2 pyrethroid colorimetric analysis.

#### b) HPLC-DAD confirmatory analysis

Quantitative analyses were carried out using Thermo Scientific™ Dionex™ Ultimate™ 3000 HPLC-DAD system (Thermofisher, Hemel Hempstead, UK) and separation achieved using an Acclaim^R^ C_18_ 120 Å (250 × 4.6 mm, Thermofisher, Hemel Hempstead, UK) column eluting with water/acetonitrile (90:10%; v/v) at a flow rate of 2 ml/min and passed through the DAD set at 275 nm. The authenticity of the detected peaks was determined by comparison of retention time, spectral extraction at 275 nm and spiking the sample with commercially available standard of the insecticide. A calibration curve of insecticide was generated by Chromeleon (Thermo Scientific™ Dionex™ Chromeleon™ 6.8 Chromatography Data System) using known amounts of the standard deltamethrin in acetonitrile injected onto the column, (0.01 – 0.4 mg/ml) for measuring the amount of insecticide on the bed net and (0.05, 0.04, 0.03, 0.02, 0.01, 0.005 mg/ml) for mosquitoes.

#### c) Measuring the amount of insecticide on the bed net

PermaNet 2.0 with the manufacturer’s claimed level of deltamethrin at 55 mg/m^2^ was used in the experiment by firstly determining the amount of deltamethrin on the net using the HPLC method as previously described [26]. Briefly, the deltamethrin concentration on the LLIN was determined at the bio-analytical laboratory at the London School of Hygiene & Tropical Medicine (LSHTM), London, UK by using HPLC-DAD. Four squares (2.5 × 2.5 cm^2^) were cut from the LLIN and each extracted using acetonitrile (1 ml) by sonication for 5 min. The supernatant was then injected into the HPLC column and the quantity of deltamethrin present was determined. Acetonitrile on its own was used as control. From this curve the amount of insecticide in the matrix was calculated. Estimated concentrations of insecticide per m^2^ were calculated from the quantities detected in each of 6.25 cm^2^ pieces.

#### d) Measuring the amount of deltamethrin on mosquitoes using HPLC-DAD

The total amount of deltamethrin extracted from each sample was measured by injecting 10 µl it onto the HPLC column and the amount of insecticide detected was expressed as mg deltamethrin/ml, and then converted to ng/mosquito. These samples were then further used in type 2-pyrethroid colorimetric test. Visual determination was compared to the sample’s deltamethrin concentration as determined by HPLC-DAD.

### Using the type 2 pyrethroid colorimetric test

#### a) Mosquito sample preparation

To confirm whether deltamethrin could be detected on either whole mosquitoes or different body parts (legs, heads, thoraxes, abdomens) using type 2 pyrethroid colorimetric test, mosquitoes were exposed to the net for 5 minutes, as described above (Fig 1). Mosquitoes were dissected using forceps that were cleaned with ethanol between insects and between body parts to avoid contamination. Pooled samples of varying numbers of mosquitoes or their body parts were used to determine what the minimum and optimal sample pool sizes were (i.e. how many mosquitoes are needed for visual detection) (Table 2).

**Table 2.** Details of the content and size of sample pools following 5-minute exposure to a treated net. These samples were used in type 2 pyrethroid colorimetric test.

| Sample pool content | Number of mosquitoes |
| --- | --- |
| Whole mosquitoes | 10, 20, 30, 40 or 50 |
| Sets of mosquito legs |  |
| Mosquito heads |  |
| Mosquito thoraces |  |
| Mosquito abdomens |  |
| Whole control mosquitoes (not exposed) |  |

#### b) Type 2 pyrethroid colorimetric test

A test invented by Kaur and Eggelte was used for the colorimetric detection of the insecticide [29]. The test entails adding potassium hydroxide (25 mM) in acetonitrile and a mixture of para nitrobenzaldehyde (6.25 mM) plus 2,3,5 triphenyltetrazolium chloride (20 mM) in acetonitrile, to each tube of homogenized mosquitoes. Glacial acetic acid (10%) in acetonitrile (solution C) was added to stop the reaction after 10 mins. The reaction produces a pink colour, and the depth of this colour gives an indication of the amount of deltamethrin present. Each test sample was compared with calibrating solutions containing known concentrations of deltamethrin (0.0001, 0.0002, 0.0004, 0.0006, 0.0008, 0.005, 0.01, 0.02, 0.03, 0.04, 0.05 mg/ml). Acetonitrile on its own was used as control blank (no colour).

Following confirmatory HPLC-DAD analysis, the same extracted samples were further tested using the screening colorimetric method, to determine how the exposure time and size of sample pools affect the results. Tubes with supernatant were left open to dry in the air for the acetonitrile to evaporate, leaving any deltamethrin residues behind. The colorimetric test was then carried out by adding solutions A, B and C, as described above.

On stopping the reaction after 10 minutes, the results were recorded with a mobile phone (Samsung Galaxy Note 8). The image was open with Microsoft Paint application and the colour of six different test spots of each sample, including control and deltamethrin standards, was quantified by recording the red, green and blue (RGB) components of each spot. An average RGB value for each sample was then calculated. These average RGB values were used to calculate ΔRGB values, which determine how much each sample differs from control, providing correlation to deltamethrin concentration [41].

## RESULTS

### Amount of insecticide on the bed net

PermaNet 2.0 was used in the experiment. We detected 55 mg/m^2^ of deltamethrin on the net using our HPLC-DAD analysis.

### Type 2-pyrethroid colorimetric test for detection of deltamethrin on mosquitoes

The colorimetric test showed that deltamethrin can be detected from whole mosquitoes and mosquito body parts (Fig. 2 and Fig. 3). Although whole mosquitoes produced the most intense colour, changes in the depth of colour could also be detected from different body parts, most often legs, heads and thoraces, but this was not consistent between different batches (Fig. 2).

**Fig. 2.**
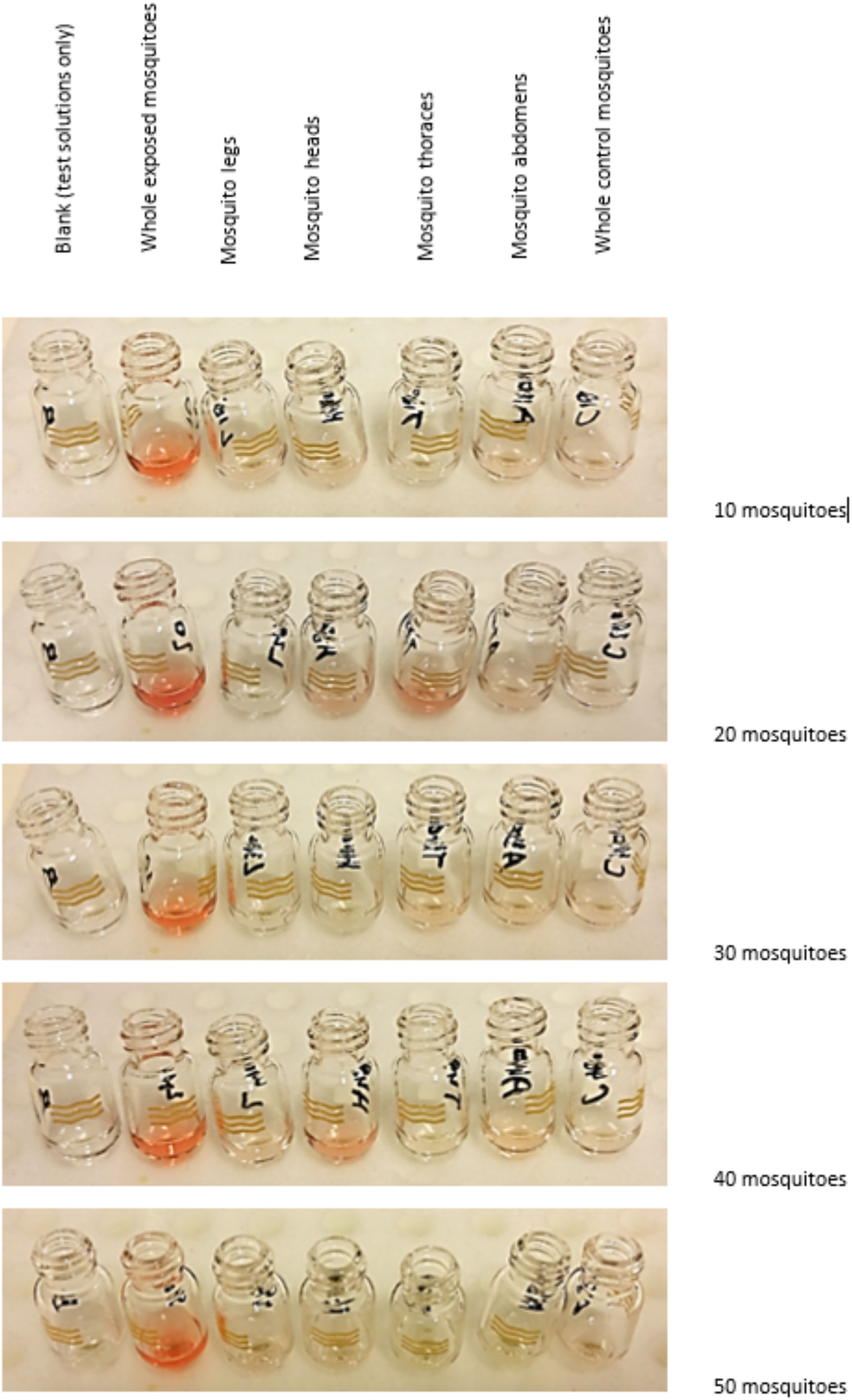
Results of the colorimetric test after 5 min exposure to PermaNet 2.0 (55 mg deltamethrin/m^2^). Whole body extracts (second vial from the left in each row) produced the strongest reaction.

**Fig. 3.**
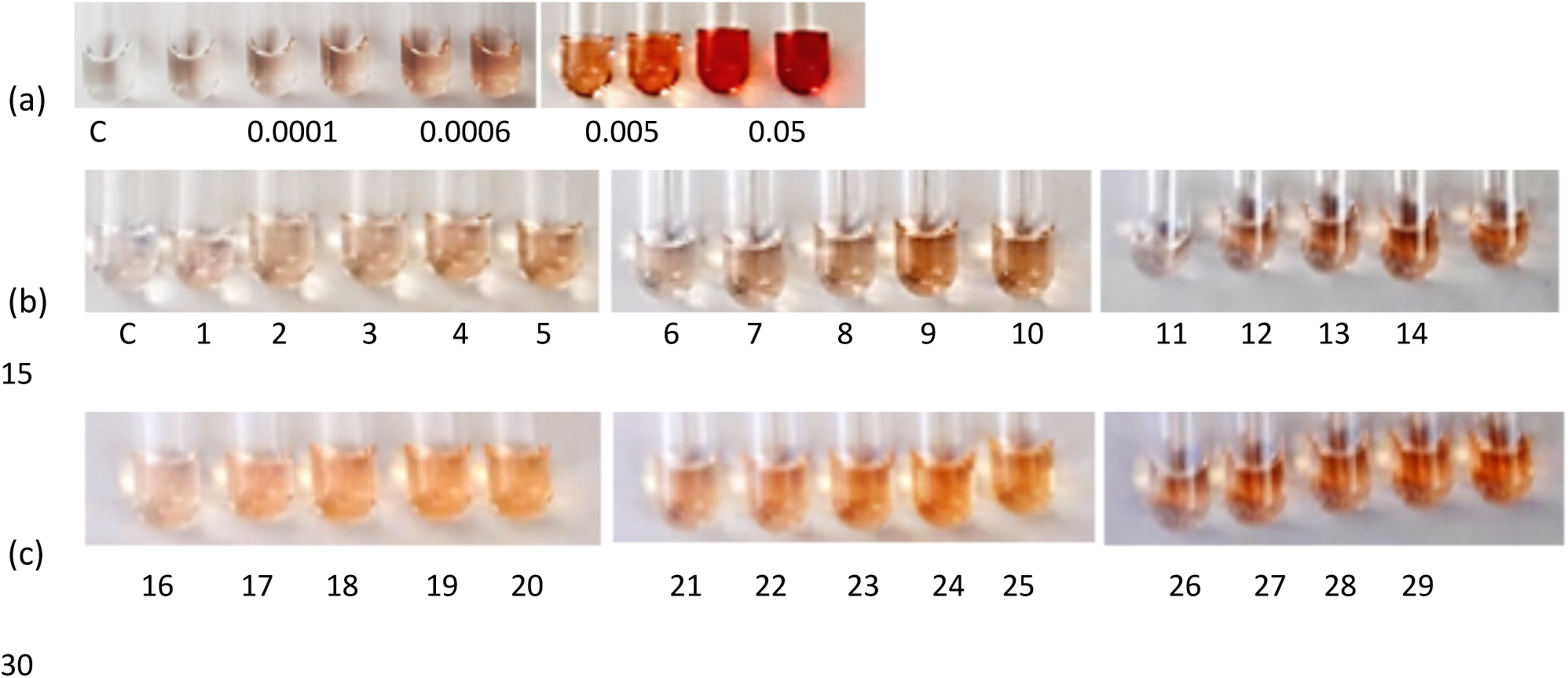
Results of the type 2-pyrethroid colorimetric test. (a) Top row left to right: deltamethrin standards at concentrations 0 (control, C), 0.0001, 0.0002, 0.0004, 0.0006, 0.0008, 0.005, 0.01, 0.02, 0.05 mg/ml. Second row left to right: control (C); samples 1-5 (0.5 min exposure), 6-10 (1 min exposure), 11-15 (1.5 min exposure). Bottom row left to right: samples 16-20 (2 min exposure), 21-25 (2.5 min exposure), 26-30 (3 min exposure).

Differences between amount of deltamethrin detected on body parts are not completely unexpected as some (e.g. legs) are more likely to come into contact with the net than others. During exposure to the net mosquitoes were at times seen trying to “crawl” through the net or were attempting to probe, which could explain stronger colouration of thorax or head samples.

### HPLC-DAD and colorimetric test experiments to measure the amount of deltamethrin on mosquitoes

Deltamethrin was measured in pools of 10 or more mosquitoes, which were exposed to a LLIN for as short as 0.5 minute (Fig. 3; samples 1 – 5 in the second row). Intensity of colour noticeably increased with the number of mosquitoes per tube and with the length of exposure time (see Table 1 for details). For example, the intensity of colour increased from sample 6 (10 mosquitoes exposed for 1 min) to sample 10 (50 mosquitoes exposed for 1 min).

Similarly, we observed an increase in the intensity of colour where pool size remained constant but the length of exposure changed: between sample 1 (10 mosquitoes, 0.5 min) to sample 6 (10 mosquitoes, 1 min), sample 11 (10 mosquitoes, 1.5 min), sample 16 (10 mosquitoes, 2 min), sample 21 (10 mosquitoes, 2.5 min), to sample 26 (10 mosquitoes, 3 min).

Samples used in the colorimetric test (as shown in Fig. 3) were first processed by HPLC-DAD. The amount of deltamethrin was measured for pools of mosquitoes (10, 20, 30, 40 or 50 mosquitoes per pool), then recalculated per mosquito (Fig. 4).

**Fig. 4.**
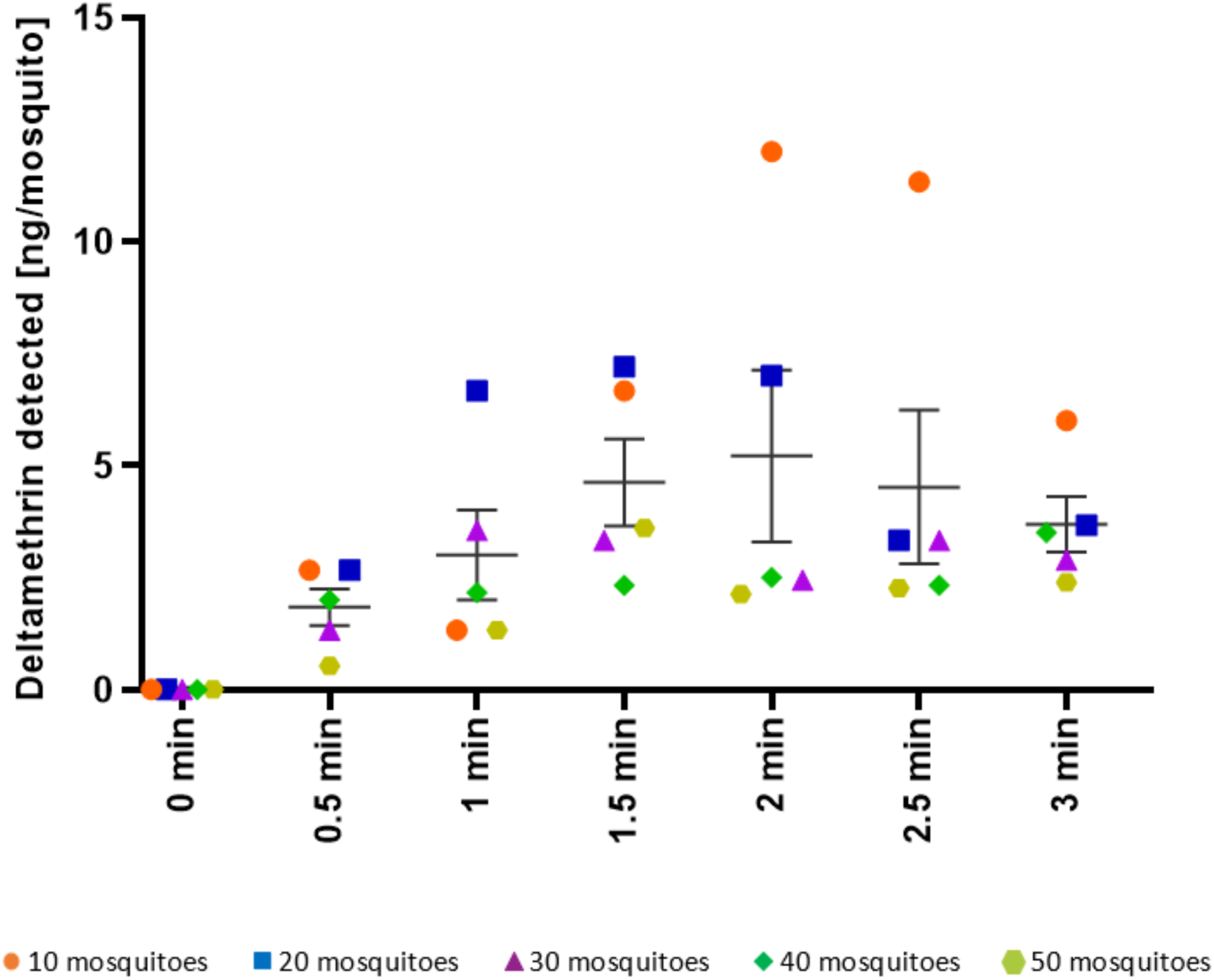
The amount of deltamethrin (ng) detected on mosquitoes exposed to PermaNet 2.0 netting for different lengths of time. Thick horizontal lines represent the mean amount of deltamethrin [ng/mosquito] for each exposure time group. Error bars represent standard error of mean.

Variation for each exposure time resulted in the significantly different (One-way ANOVA, F_6,28_ = 2.576, *p* = 0.0409) amounts of insecticide measured. There was an increasing trend with more deltamethrin present on mosquitoes that were exposed to the net for longer, between 0.5 minute exposure (1.8 ng/mosquito) and 2.0 minute exposure (5.2 ng/mosquito) (*p = 0*.*0110*); no significant difference was observed in the amount of deltamethrin detected on mosquitoes that were exposed to the net between 2.0 minutes and 3 minutes (3.7 ng/mosquito).

There was good correlation between the amount of deltamethrin measured by HPLC-DAD and intensity of colour obtained using the colorimetric test for majority of samples. Using digital images and image analysis process with RGB values, it is possible to detect changes in colour intensity and therefore estimate how much insecticide mosquitoes pick from treated surfaces (Fig. 5).

**Fig. 5.**
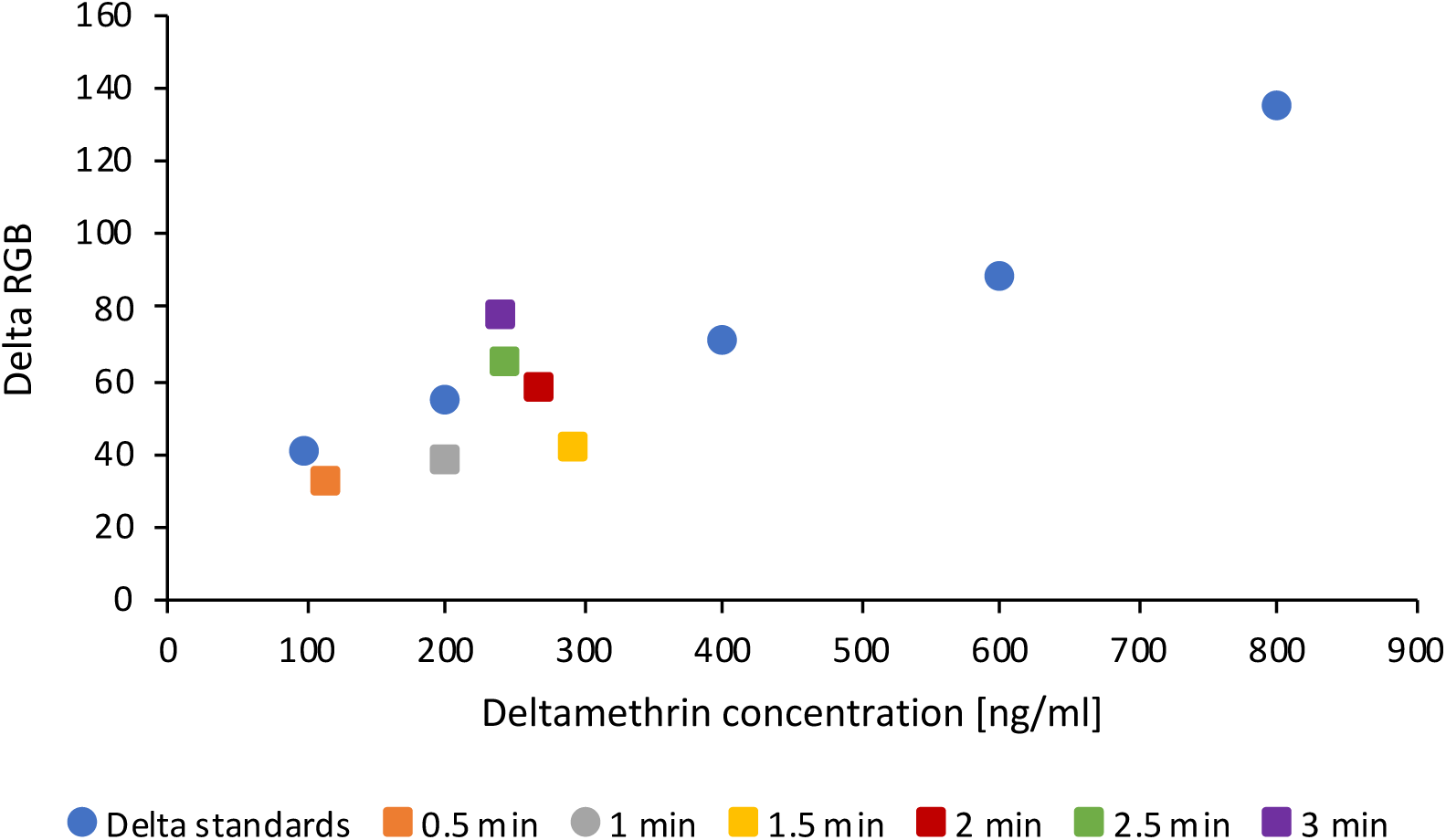
Relationship between intensity of colour produced in comparison to control, expressed as ΔRGB, and deltamethrin concentration (ng/ml) detected in samples of known concentration (delta standards) and mosquito samples exposed to PermaNet 2.0 netting for different lengths of time. Concentration of deltamethrin for mosquito samples is given in ng/ml and not in ng of insecticide per mosquito.

## DISCUSSION

While pyrethroids are currently the only insecticide class used on all ITNs, next-generation LLINs treated with a combination of pyrethroids and synergists or non-pyrethroid insecticides are currently under evaluation [2, 42]. Alternatively, pyrethroid LLINs can be used in combination with non-pyrethroid IRS. Mixtures of active ingredients, and combinations of interventions have been proposed as available strategies for insecticide resistance management [43]. Various methods can be used to determine the quantity of insecticides present on the ITNs or sprayed walls for the purposes of quality control, operational monitoring of spraying operations or to monitor degradation of insecticides on ITNs over time [26, 27, 32, 33, 38, 44, 45]. As the new combined tools are introduced, detection of active ingredients that actually come into contact with mosquitoes and cause the greatest mortality might be additionally used in assessing the effectiveness of interventions. Furthermore, measuring the amount of insecticide mosquitoes pick up when they come in contact with treated surfaces will inform future decisions on the doses of active ingredients used in vector control tools and be a part of insecticide resistance management.

Pyrethroids need to penetrate through the mosquito cuticle to reach their target sites in the nervous system. The insecticides point of entry is either through mosquito tarsi when the insects land on the treated surfaces, or through mosquito body if they collide with the net [8, 46]. Mosquito – LLIN interactions have been characterized using infrared video tracking, showing that susceptible mosquitoes made between 11.0 and 57.1 seconds of contact with a LLIN during the initial 10-minute period of most intense mosquito activity around the net [8]. Mean time spent on deltamethrin-treated net by a susceptible mosquito, causing knockdown and death, was measured to be 70.1 seconds, with the minimum required to cause knockdown just 0.4 seconds [24]. The exposure time in our experiments was chosen accordingly, starting at 30 seconds, whereas the longest exposure time (3 minutes) was the same as that used in WHO standard method for LLIN evaluation [47].

Little is known about the actual amount of insecticide that adheres to the mosquitoes after contact with treated surfaces. Previous studies have measured the amount of DDT on mosquitoes which entered sprayed huts (sprayed with 200 µg/cm^2^ active ingredient) using gas chromatography [18]. The amount of DDT on dead *An. gambiae* and *An. funestus* was in the range of 7–20 ng/mosquito, whereas much lower levels of DDT (around 1.5 ng/mosquito) were found on surviving mosquitoes. Another study measured the amount of dieldrin picked up by *Culex quinquefasciatus* during the exposure in standard WHO bioassay tubes, using different concentrations on papers and different exposure times [17]. The authors concluded that pick-up of insecticide is a linear function of both the concentration and exposure time. It appears that at least some insecticide becomes internalised rapidly after exposure. When deltamethrin was topically applied to mosquito legs, about 5% of the initial applied amount could be detected in the mosquito body after 15-minute exposure (i.e. 0.048 ng/ susceptible mosquito body) [48]. When mosquitoes fed through a radio-labeled permethrin net the insecticide was shown to reach the midgut and was detected in the blood meal within an hour after feeding using a scintillation counter [13]. Most recently, liquid chromatography tandem mass spectrometry was used to measure insecticide load per *An. gambiae* female mosquito following a standard WHO susceptibility test with 0.05% deltamethrin paper, detecting 34 – 52 pg of deltamethrin per mosquito [19].

Through this study we have established that the amount of deltamethrin detected per mosquito using HPLC-DAD increased between 0.5 – 2 minute exposure and then levelled off during longer exposure times (Fig. 3), which agrees with observations by Pennell *et al* of the amount of insecticide dieldrin present on the exterior of mosquitoes as opposed to “internal” amount which increased with increasing exposure time and concentration [17]. The type of insecticide used, its penetration through the cuticle, possible accumulation in the hemolymph and internal organs, its mode of action and its metabolism within the insect will all determine what happens to insecticides after the initial contact [10]. Pyrethroids are known to associate with hemolymph carrier proteins and with lipids [9]. The mosquito samples used in our experiments for HPLC-DAD analysis were pulverised by grinding and prolonged sonication and homogenization before extraction with acetonitrile which dissociates and dissolves the deltamethrin. The lower limit of detection of deltamethrin on HPLC-DAD was 19.2 ng/ml which is not sensitive enough to measure the exact amount of insecticide per mosquito. Hence, pools of ten to fifty mosquitoes were used to measure the collective amount of adhered insecticide, then expressed per mosquito.

Variability between individual insects has been observed in nature, caused by differences in mosquito age, blood-feeding status, the presence of different resistance mechanisms and any fitness costs they might incur, and differences in mosquito behaviour in relation to LLINs and frequency of exposure [49]. Some resistance genes, such as *kdr*, are only weakly related to survival and their presence also means that mosquitoes are less repelled by pyrethroids, so they tend to acquire a higher dose of insecticide which can be lethal [20]. Yet susceptible mosquitoes might be irritated by pyrethroids before they are killed, which could result in large variation between individuals and the amounts of insecticide they pick up – hence there can really be no one “discriminating dose”. Some susceptible mosquitoes might pick up only a small amount of insecticide and survive, while some resistant mosquitoes might pick up significantly more and die. This can manifest as variation between replicates of WHO bioassays, and can affect to what extent LLINs control local mosquito populations.

This investigation further demonstrated that colorimetric test is easy to use, providing results within 15 mins detecting the presence of deltamethrin not only on ITNs [31] and sprayed walls [32] but also on mosquitoes which in this investigation came into contact with treated PermaNet 2.0 netting. A change in the depth of colour was detected in all samples, including the sample with the fewest pooled mosquitoes (10) and the shortest exposure time (30 seconds). Moreover, pyrethroids were also detected in pools of different mosquito body parts, but with less consistency as the amounts of insecticide were that much smaller. It has been shown that mosquitoes obtain particles across their entire body in a standard 3-minute WHO cone bioassay and that particles can be transferred to their legs even following short contact periods [46]. Using only parts of collected mosquitoes could be advantageous during field work when the rest of the mosquito is required for other tests (eg. Blood meal analysis, or to test for the presence of *P. falciparum* infection) but would need to be tested in larger pools.

Overall there was good correlation between the two analysis methods used. We have also shown for the first time that the amount of deltamethrin mosquitoes acquire after coming into contact with a LLIN is in the range of up to 10 ng/mosquito. This information can be further used to improve dosing of insecticides on treated nets or sprayed walls and can be used in future studies of other compounds used for vector control.

## DECLARATIONS

### Ethics approval and consent to participate

Not applicable.

### Consent for publication

Not applicable.

### Availability of data and material

The datasets used and/or analysed during the current study are available from the corresponding author on the reasonable request.

### Authors’ contributions

MK and HK conceived and developed the study design with contributions from JL. MK reared the mosquitoes, exposed them to deltamethrin and prepared samples for further analysis. HK carried out HPLC-DAD analysis. MK and HK performed the colorimetric test. MK analysed the data with HKs assistance. MK wrote the first draft of the manuscript with inputs from HK and JL. All authors reviewed the manuscript, provided comments and approved the final manuscript.

## Acknowledgements

Not applicable.

